# Polyfunctional tumor-reactive T cells are effectively expanded from non-small cell lung cancers, and correlate with an immune-engaged T cell profile

**DOI:** 10.1101/426221

**Authors:** Rosa de Groot, Marleen M. van Loenen, Aurélie Guislain, Benoit P. Nicolet, Julian J. Freen-van Heeren, Onno Verhagen, Michel M. van den Heuvel, Jeroen de Jong, Patrick Burger, C. Ellen van der Schoot, Robbert M. Spaapen, Derk Amsen, John B. A. G. Haanen, Kim Monkhorst, Koen J. Hartemink, Monika C. Wolkers

## Abstract

Non-small cell lung cancer (NSCLC) is the second most prevalent type of cancer. With the current treatment regimens, the mortality rate remains high. Therefore, better therapeutic approaches are necessary. NSCLCs generally possess many genetic mutations and are well infiltrated by T cells (TIL), making TIL therapy an attractive option. Here we show that T cells from treatment naive, stage I-IVa NSCLC tumors can effectively be isolated and expanded, with similar efficiency as from normal lung tissue. Importantly, 76% (13/17) of tested TIL products isolates from NSCLC lesions exhibited clear reactivity against primary tumor digests, with 0.5%-30% of T cells producing the inflammatory cytokine Interferon (IFN)-γ. Both CD4^+^ and CD8^+^ T cells displayed tumor reactivity. The cytokine production correlated well with CD137 and CD40L expression. Furthermore, almost half (7/17) of the TIL products contained polyfunctional T cells that produced Tumor Necrosis Factor (TNF)-α and/or IL-2 in addition to IFN-γ, a hallmark of effective immune responses. Tumor-reactivity in the TIL products correlated with high percentages of CD103^+^CD69^+^CD8^+^ T cell infiltrates in the tumor lesions, with PD-1^hi^CD4^+^ T cells, and with FoxP3^+^CD25^+^CD4^+^ regulatory T cell infiltrates, suggesting that the composition of T cell infiltrates may predict the level of tumor reactivity. In conclusion, the effective generation of tumor-reactive and polyfunctional TIL products implies that TIL therapy will be a successful treatment regimen for NSCLC patients.

## INTRODUCTION

Non-small cell lung cancer (NSCLC) is one of the most prevalent types of cancer worldwide. With the current treatment regimens, the 5-year survival rate of patients suffering from NSCLC is still limited. Because of the striking effects of immunotherapy in melanoma (1–4), recent efforts are directed towards treatment with immunotherapy for NSCLC patients. NSCLC are considered immunogenic, because they often contain high numbers of somatic mutations which is induced by smoking tobacco (5). Furthermore, they are well infiltrated by T cells (6–8). The functionality of these tumor infiltrated T lymphocytes (TILs) has, however, not been well documented. Treatment with anti-PD-1 is currently standard of care for stage IV NSCLC patients with distant metastasis. This therapy resulted in tumor regression in 17-30% of the patients (9–13). The majority of patients with NSCLC thus does not benefit from immune checkpoint blockade. Tumors in these patients utilize different immunosuppressive pathways (14) that need to be overcome in order for immunotherapy to work. Therefore, additional immunotherapy interventions should be explored for the treatment of NSCLC patients, possibly in combination with the existing anti-PD-1 therapy.

One therapeutic opportunity for intervention is the infusion of TILs. Adoptive transfer of *ex vivo* expanded TILs has proven highly effective for stage IV melanoma patients (15), with impressive 50% overall response rates in pretreated patients (1, 2). Of these melanoma patients, 10-20% experience durable complete remission (1, 2).

Tumor-reactive T cells were already detected in the mid 1990s in NSCLC lesions (16–18), and a clinical effect of TIL therapy for stage IV NSCLC patients has been reported, albeit with very minor improvements in survival (19). Since that time, the treatment regimen substantially improved, by speeding up the protocols to culture and expand TILs from tumor lesions (1, 2), and by pre-conditioning the patient with non-myeloablative chemotherapy (20) that allowed for the above-mentioned success rates in melanoma patients. Therefore, the efficacy to grow tumor-reactive TIL products from NSCLC lesions should be re-assessed, both in terms of cell expansion and the presence of cytokine-producing TILs in response to tumors. Furthermore, it is yet to be determined whether a specific T cell profile in tumor lesions correlates with the level of tumor reactivity of expanded TIL products.

Here, we show that most TIL products contain tumor-reactive T cells. In particular TIL products with high tumor reactivity are polyfunctional. Furthermore, tumor reactivity of the expanded TIL product correlated with the composition of the T cell compartment in the tumor lesions. We conclude that the generation of NSCLC-specific TIL products for therapeutic purposes is feasible and should be reconsidered for clinical application.

## Materials and methods

### Patient cohort and study design

Between June 2015 and June 2017, 25 treatment-naive NSCLC patients were included in this study. Samples from 2 patients were excluded because of logistic issues. Table 1 depicts the patient characteristics of the remaining 23 donors. The cohort consisted of 10 male and 13 female donors between the age of 38 and 79 years (average 66,1 years) with clinical stage Ia-IVa according to the TNM7 staging system for NSCLC, based on tumor size, nodal involvement and level of metastasis. All but two patients had a history of smoking.

**Table 1:** Patient characteristics

| **Donor**<br>**(N=25)** | **Clinical stage**<br>**(TNM7\*)** | **Gender**<br>**(M/F)** | **Age at surgery**<br>**(Years)** | **Smoking history**<br>**(Cigarettes/day)** |
| --- | --- | --- | --- | --- |
| **1** | IIA | M | 71 | no (never) |
| **2** | IIA | M | 53 | yes (15/d) |
| **3** | IIB | M | 77 | yes (3/d) |
| **4** | 0-IA1 | F | 67 | yes (8/d) |
| **5** | IIB | F | 57 | yes (10/d) |
| **6** | IA1 | F | 59 | yes (3/d) |
| **7** | IIA | F | 61 | yes (15/d) |
| **8** | IVA | M | 64 | yes (25/d) |
| **9** | IIA | M | 60 | yes (20/d) |
| **10** | IIB | F | 56 | yes (30/d) |
| **11** | IB | M | 59 | yes (20/d) |
| **12** | IIB | F | 70 | yes (20/d) |
| **13** | IA1 | F | 58 | yes (8/d) |
| **14** | IA2 | F | 38 | yes (20/d) |
| **15** | IA2 | F | 81 | no (never) |
| **16** | IIB | F | 63 | yes 20/d) |
| **17** | IB | F | 69 | yes (10/d) |
| **18** | IB | F | 56 | yes (6/d) |
| **19** | IB | F | 79 | yes (5/d) |
| **20** | IIIA | M | 73 | yes (30/d) |
| **21** | IA3 | F | 51 | yes (8/d) |
| **22** | IIIB | M | 61 | yes (15/d) |
| **23** | *excluded* |  |  |  |
| **24** | *excluded* |  |  |  |
| **25** | IA3 | M | 69 | yes (20/d) |

*\*American Joint committee on Cancer: Lung Cancer Staging, 7<sup>th</sup> edition, 2009* *M=Male; F=Female; d=day*

The study protocol was designed to define the phenotype of TILs isolated from treatment naive NSCLC compared to non-tumorous lung tissue (N=16), to explore the capacity to expand TILs (N=17) and to measure the presence of tumor-reactive T cells within TIL cultures (N=17). The study was performed according to the Declaration of Helsinki (seventh revision, 2013), and executed with consent of the Institutional Review Board of the Netherlands Cancer Institute-Antoni van Leeuwenhoek Hospital (NKI-AvL), Amsterdam, the Netherlands. Donor tissues were obtained directly after surgery, transported at room temperature in RPMI medium 1640 (Gibco) containing 50ug/ml gentamycin (Sigma-Aldrich), 12,5ug/ml Fungizone (Amphotericin B, Gibco) and 20% fetal bovine serum (FBS) (Bodego), weighed and processed within four hours. Distal healthy lung tissue as defined by the pathologist was taken as far away as possible from the tumor lesion.

Tumor stage and differentiation, and weight of the obtained tissues for this study are shown in Table 2. Tumor size ranged from T1a-T4 according to the TNM7 system, and specified as adenocarcinoma (n=14), squamous carcinoma (n=5) and as NSCLC not otherwise specified (NOS, n=4). On average 1420 mg tumor tissue was obtained, ranging from 23-9590 mg (Table 2).

**Table 2:**
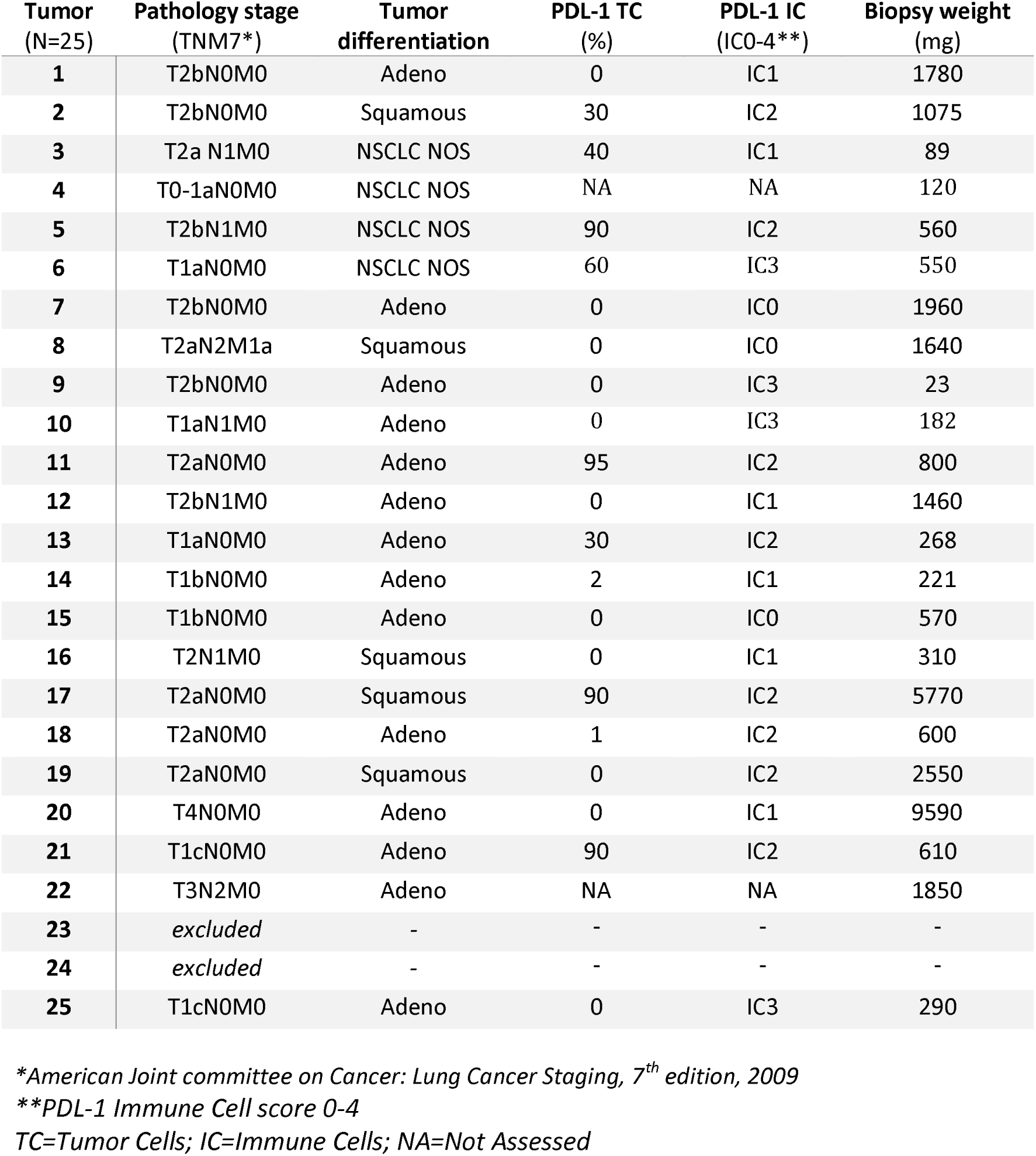
Tumor characteristics

### Tissue digestion

The first five donors were used to optimize the digestion protocol for the highest yield of live CD45^+^ cells in single cell digests. Briefly, freshly isolated tumorous and non-tumorous lung tissue were finely chopped, and incubated for 45 min shaking at 37°C in IMDM (Gibco) containing 30IU/ml collagenase IV (Worthington), 12,5μg/ml DNAse (Roche), and 1% FBS. The digest was pelleted at 360g for 10 min, and resuspended in FACS buffer (PBS containing 2% FCS and 2mM EDTA). Digest was filtered first through a tea mesh and then over a 100um filter. After red blood cell lysis for 15 min at 4°C with 155 mM NH_4_Cl, 10 mM KHCO_3_, 0.1 mM EDTA (pH 7.4), live and dead cells were manually counted with trypan blue solution (Sigma) on the hemocytometer. From six donors, the tumor tissue was divided in three regions. A sample from each region was separately processed for TIL culture. The remaining tissue was combined for the total tumor digest. After digestion, 1-2×10^6^ live cells were used for flow cytometry analysis, and 1-3×10^6^ live cells were used for TIL cultures. The remaining digest was cryo-preserved until further use.

### T cell expansion

TIL cultures and T cell cultures from distal lung tissue digests were performed as previously described for melanoma-derived TILs (1, 2). Briefly, 2-3 wells containing 0,5-1*10^6^ live cells from the tissue digest were cultured for 10-13 days in 24 wells plates in 20/80 T-cell mixed media (Miltenyi) containing 5% human serum (HS) (Sanquin), 5% FBS, 50μg/ml gentamycin, 1,25 μg/ml fungizone, and 6000 IU human recombinant (hr) IL-2 (Proleukin, Novartis) (pre-Rapid Expansion Phase; pre-REP).

Medium was refreshed on days 7, 9 and 11. Wells were split when a monolayer of cells was observed in the entire well. When more than 30% of the cells stopped dividing (determined as rounding up of cells), cells were harvested, counted and prepared for an additional culture period of 10-13 days (REP). 1-3 times 2×10^5^ live cells/well (1-3 wells/donor) were co-cultured with 5-10×10^6^ irradiated PBMCs pooled from 10 healthy donors in 24 wells, 30ng/ml anti-CD3 antibody (OKT-3) (Miltenyi Biotec) and 3000IU/ml IL-2. Typically, cells were passaged at day 5, 7, 9 and 11 and harvested, washed and counted on day 10-13, based on visual assessment of the T cell proliferation state as above. Cells were either tested immediately for reactivity, or cryo-preserved in IMDM containing 10% DMSO (Corning) and 30% FBS until further use.

### T cell activation

After thawing and live cell counting as described above, pre-REP and REP T cells were pre-stained in FACS buffer with anti-CD4 BUV496, anti-CD8 BUV805 (BD Biosciences) for 30 min at 4°C. Cells were washed twice, one time in FACS buffer and then in 20/80 T-cell mixed media (Miltenyi). 100.000 live T cells were co-cultured with 200.000 live tumor digest cells or normal lung digest cells for 6h at 37°C, were stimulated with 10ng/ml PMA and 1ug/ml Ionomycin (SigmaAldrich), or cultured with T-cell mixed media alone. Brefeldin A (Invitrogen) was added after 30 min of co-culture.

### Flow cytometry

For ex vivo analysis of T cells in the tissue digests, cells were washed in FACS buffer, and then stained in FACS-buffer for 30 minutes at 4°C with the following antibodies: anti-CD3 PerCp-Cy5.5, anti-CD279 FITC, anti-CD56 BV605, anti-CD27 BV510, anti-CD127 BV421, anti-CD103 PE-Cy7 and anti-CD25 PE (all from Biolegend), and with anti-CD8 BUV805, anti-CD45RA BUV737, anti-CD4 BUV496, and anti-CD69 BUV395 (from BD Biosciences). Alternatively cells were stained for anti-Lag3 PE (BD Biosciences), and anti-TIGIT Alexa 647 (BD Biosciences). Live/dead fixable near IR APC-Cy7 (Invitrogen) was included for dead cell exclusion. Cells were washed twice with FACS buffer and fixed for 30 minutes with the Perm/Fix Foxp3 staining kit (Invitrogen) according to the manufacturer’s protocol using anti-Foxp3 Alexa647 (Biolegend). Cells were resuspended in FACS buffer and passed through a 70μm single cell filter prior to flow cytometry analysis (LSR fortessa, BD biosciences). The first six donors were used to optimize the staining panel and cytometer settings, by antibody titration and the use of single stains in combination with the minus one fluorochrome method (MOF).

T cells from pre-REP or REP cultures were stained prior to T cell activation with anti-CD4 BUV496, anti-CD8 BUV805 as described above. After the full activation protocol, cells were washed twice with FACS buffer and stained with anti-CD154 BV510 and anti-CD279 BV421 (Biolegend) and Live/dead fixable near IR APC-Cy7 in FACS buffer for 30 minutes at 4°C, and washed twice in PBS. Cells were then fixed and stained with anti-CD137 PE-Cy7, anti-IFN-γ PE, anti-TNF-α Alexa488 and anti-IL-2 APC (Biolegend) with the Cytofix/CytoPerm staining kit (BD Biosciences) according to the manufacturer’s protocol. Anti-CD107 Alexa700 (BD Biosciences) was included during the co-culture with tumor cells. Cells were washed in FACS buffer and passed through a 70uM single cell filter prior to acquisition with the LSR fortessa (BD). Flow cytometry settings were defined for each patient material using single stainings for each antibody. To each flow cytometry analysis of patient material, a standardized sample of PBMCs pooled from ten healthy donors that was cryo preserved prior to the start of the study was included. Data analysis was performed with Flowjow Star 10.1.

### Immunohistochemistry on tumor tissue

PD-L1 expression on formalin-fixed, paraffin-embedded tissue samples was assessed with immunohistochemistry using the monoclonal antibody 22C3 from Agilent on a Ventana Benchmark Ultra. At least 100 neoplastic cells were scored for membranous staining. Tumor cell PD-L1 scores were described as percentages of total tumor cells. ICs are scored as the proportion of tumor area that is occupied by PD-L1 staining of immune cells of any intensity and categorized in IC0-4 representing the following cutoffs; 0%, 0 - 5%, 5 - 10% and >10%.

### Clonal T cell analysis

For clonal T cell analysis, CD3^+^CD8^+^ T cells and CD3^+^CD4^+^ T cells (excluding CD25^hi^CD127^−^ T cells to exclude Tregs) from tumor digests and from the corresponding REP TIL were FACS-sorted. Cells were washed with PBS, and cell pellets snapfrozen in liquid nitrogen were stored at −30 °C. DNA extraction was performed with NucleoSpin Tissue XS (Macherey Nagel) according to the manufacturer’s instructions. Complete TCR beta rearrangements were amplified in multiplex PCR with sets of Vbeta- and J beta-specific primers, respectively. In a nested PCR (20 cycli) barcodes and the Ion Torrent specific adaptors A en P1 were added to the PCR products. The DNA library was sequenced on the Ion S5 sequencer according to manufacturer’s instruction. Files containing the Unprocessed sequencing reads (fastq) were analysed with mixcr version 3.0.5 (21) using “TCR output only” option. Txt output was then imported in R (3.5.1) under R Studio (1.1.453) and further analyzed using tcR version 2.2.3 (22). Clone frequency and shared repertoire analysis on nucleic-acid sequences was analyzed with tcR. CD4 and CD8 clones in each donors were filtered using tcR and plotted in bar graphs made in R using ggplot 2 version 3.0.0.

### Stasticial analysis

Statistical analyses were performed with Graphpad Prism 7.2. Compiled data are shown as paired data points for each patient, or as single data points with box and whiskers showing maximum, 75th percentile, median, 25th percentile, minimum and mean, unless otherwise indicated in the legend.

Overall significance of differences were calculated with repeated measurement paired one-way ANOVA test with Tukey’s post hoc test comparing columns. If differences were confirmed (p<0,05), significance between two data points was calculated using paired Student’s t test, with the p value cut-offs of *= p<0.05; **=p<0.01; and ***=p<0.001. If Student’s t test showed p values ≥0.05, p value marking was omitted in panels. Correlations were calculated using Pearson’s correlation in combination with linear regression.

## RESULTS

### High yield of lymphoid cells isolated from NSCLC tumor lesions

We first determined the efficacy of isolating TILs from NSCLC lesions that onderwent lobectomy. 23 patients from treatment-naive stage Ib-IVa NSCLC patients suffering from non-squamouse (n=14), squamous (n=5), or from NSCLC not otherwise specified (n=4) were included in this study (Table 1). To evaluate the TIL isolation and expansion procedure from the tumor, we also isolated normal lung tissue from the same patients that was harvested as far away as possible from the tumor lesion.

From the tumor digests, we obtained on average 33.6×10^3^ viable cells/mg tissue, which was comparable to the yield from normal lung tissue digests, with on average 51.2×10^3^ viable cells/mg tissue (Fig. 1A). In line with previous studies (7, 8, 23), we detected high numbers of CD3^+^CD56^−^ T cells cells in both tumor and normal lung tissues, with 23.7±16.9% and 15.5±13.1% of live cells in the respective tissue digests. This translated into on average 17.8×10^3^ T cells/mg lung tissue and 9.8×10^3^ T cells/mg tumor tissue (Fig. 1B-D). Of note, sufficient cell numbers for TIL expansion and phenotypic analysis could be isolated even from tumors of <1 cm^3^ (Table 2).

**Figure 1:**
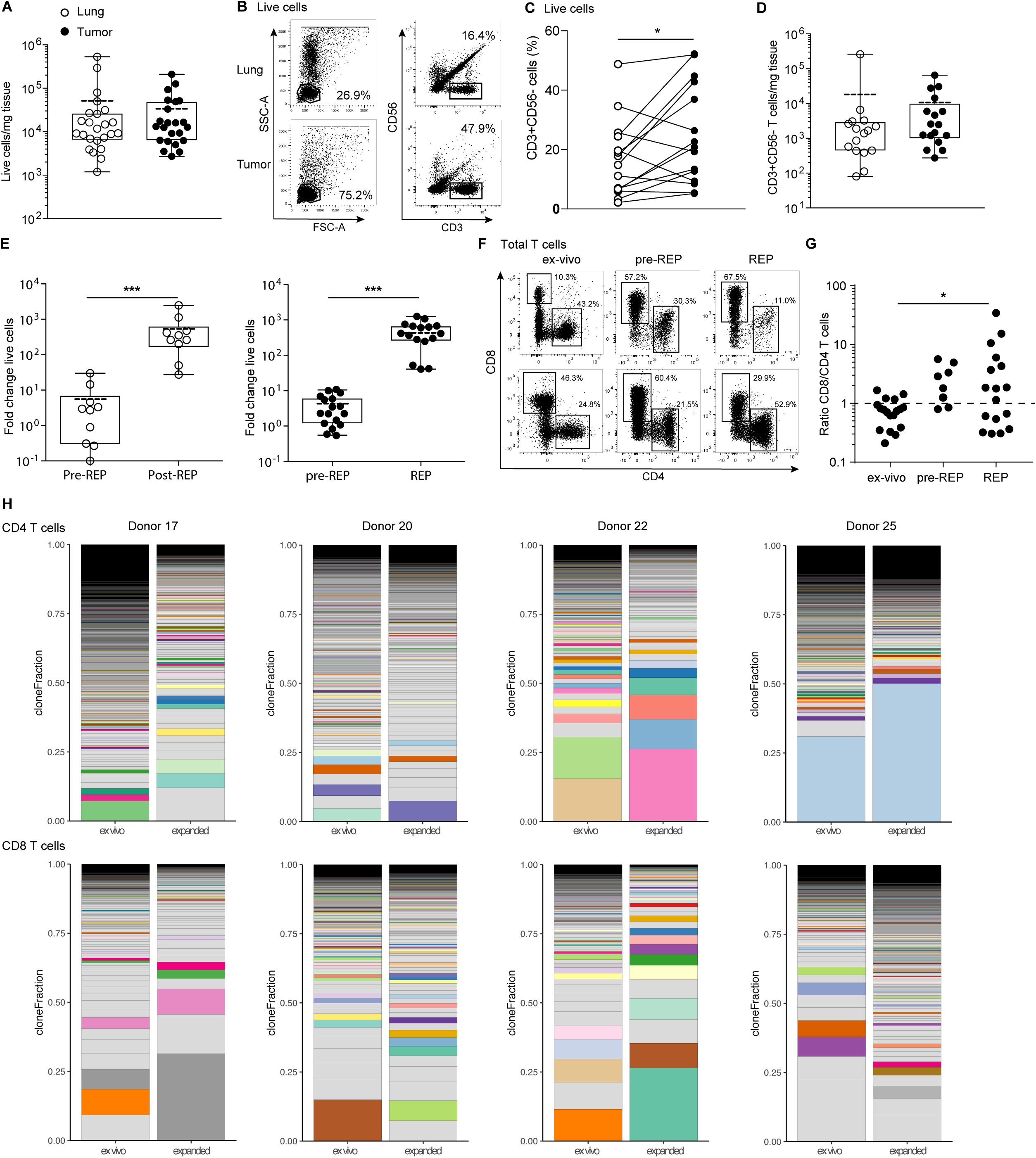
Effective isolation and expansion of TILs from NSCLC-derived lesions. Single cell suspensions were obtained from distal lung tissue and tumor sections of 23 donors directly after resection. **(A)** Numbers of life cells/mg tissue. Dead cells were excluded by trypan blue. Each dot represents one donor, box and whisker plots depict minimum, 25^th^ pct, median, 75^th^ pct and maximum values. Mean is depicted by the dotted line. **(B)** Gating strategy of CD56^−^/CD3^+^ T cells and lymphocytes, and **(C)** percentage of CD3^+^ cells within the lymphocyte population in tumor and lung, to calculate **(D)** the numbers of T cells/mg tissue by flow cytometry, depicted as in (A). **(E)** Of 9 normal lung tissues (left panel), and of 17 tumor tissues (right panel), single cell suspensions were cultured for 10-13 days with 6000IU/ml IL-2 (pre-REP), followed by 2 weeks of culture with 30ng/ml anti-CD3, feeder mix, and 3000IU/ml IL-2 (REP). The fold change of total cell count was determined from the number of cells used as input. **(F, G)** The ratio of CD8^+^ over CD4^+^ T cells of the CD3^+^ T cells was determined by flow cytometry in tumor digests directly ex vivo, and after pre-REP and REP cultures. [Paired student’s T test; * p<0.05; *** p<0.001. If no indication, p≥0.05]. **(H)** T cell clonality was determined by CDR3 sequencing of the TCRb chain of FACS-sorted CD4^+^ and CD8^+^ T cells from tumor digest and from corresponding expanded TILs (CD25^hi^CD127^hi^ CD4^+^ T cells containing the Treg population were excluded from this analysis). Relative individual T cell clone frequency ordered from the largest (bottom) to the smallest (top). Color-coded pairs represent shared TCR sequences, light-grey bars depict non-shared TCR sequences

### Effective TIL expansion from NSCLC tumors

The expansion efficiency of TILs highly depends on the tumor type and the culture protocol (1, 2). We here determined the efficiency of TIL expansion from 17 tumor digests with the rapid expansion protocol (REP) established for melanoma TILs (1, 2). For comparison, we included 9 normal lung tissue digests. Tumor and lung tissue digests were cultured for 10-13 days in the presence of IL-2 (pre-REP), followed by a 10-13 day restimulation with the α-CD3 antibody OKT-3 and IL-2 (REP). During pre-REP, CD3^+^ T cells from tumor and lung digest expanded on average 4- and 6-fold, respectively (Fig. 1E). REP cultures expanded CD3^+^ T cells from the tumor 470-fold and from the lung 570-fold, resulting in an overall expansion of on average 2000-fold for tumor TILs, with a minimum of 540-fold, and 3000-fold lung tissue derived T cells (Fig. 1E). Thus, even though T cells from normal lung tissue may expand slightly better than those from the tumor, the overall efficiency of expanding NSCLC-derived T cells was comparable to recently reported expansions of melanoma-derived TILs (1, 2).

We then measured the percentages of CD4^+^ and CD8^+^ T cells in the pre-REP and REP cultures from tumor digests. CD4^+^ T cells increased from 14.3± 9.6% ex vivo of total live cells to 43.3±26.5% after REP, and CD8^+^ T cells increased from 10.3±8.3% to 53.4±27.0% (Fig. 1F). CD8^+^ T cells expanded generally better than CD4^+^ T cells in these culture conditions, as revealed by the changes over time in the CD8^+^/CD4^+^ T cell ratio (Fig. 1G). The rate of T cell expansion did not correlate with the PD-L1 expression on tumor cells or immune infiltrates, or with the T cell differentiation status (naïve/effector/memory) within the tumor tissue (Supplemental Fig. 1).

We next investigated the phenotypic alterations in T cell composition of the expanded TIL products. TIL expansion resulted in a drop of FOXP3-expressing CD4^+^ to <0.1% of the CD4 T cells (Supplemental Fig. 2A). Furthermore, even though LAG-3 and TIGIT expression increased upon T cell cultures, the expression of PD-1 expression substantially diminished upon T cell expansion (Supplemental Fig. 2B). We also interrogated whether and how the T cell clonality altered upon T cell expansion.

CDR3 nucleic acid sequence analysis of the TCRβ chain revealed that both CD4^+^ and CD8^+^ T cells are in general polyclonal in NSCLC tumor lesions, and that this polyclonality is maintained upon expansion (Fig. 1H). Furthermore, tumor-derived and expanded CD4^+^ and CD8^+^ T cells showed overlapping T cell clones, a feature that was more pronounced in CD8^+^ T cells (Fig. 1H, Supplemental Fig. 3). In conclusion, both CD4^+^ and CD8^+^ T cells from NSCLC expanded well with the REP culture conditions, and maintained their polyclonal profile.

### The majority of TIL products contain tumor-specific T cells

We next examined the tumor reactivity of expanded TILs. As a read-out, we measured the production of the pro-inflammatory cytokine IFN-γ by expanded TILs upon co-culture with autologous tumor digest. This production was compared to co-culture with normal lung tissue digest, or with medium alone. To distinguish the expanded TILs from the high T cell infiltrates within the tissue digests, we pre-stained the expanded TILs with CD4 and CD8 antibodies prior to co-culture.

IFN_-_γ producing T cells in response to tumor tissue were detected in 16/17 tumor digests (Fig. 2A,B). On average, 6.3±7.8% of the T cells from the TIL products produced IFN-γ when co-cultured with tumor digests (Fig. 2A,B). We detected similar percentages of IFN-γ-producing T cells within the total CD4^+^ and CD8^+^ T cell population, with 5.8±6.3% and 6.5±9.4% of each population, respectively (Fig. 2A,B).

**Figure 2:**
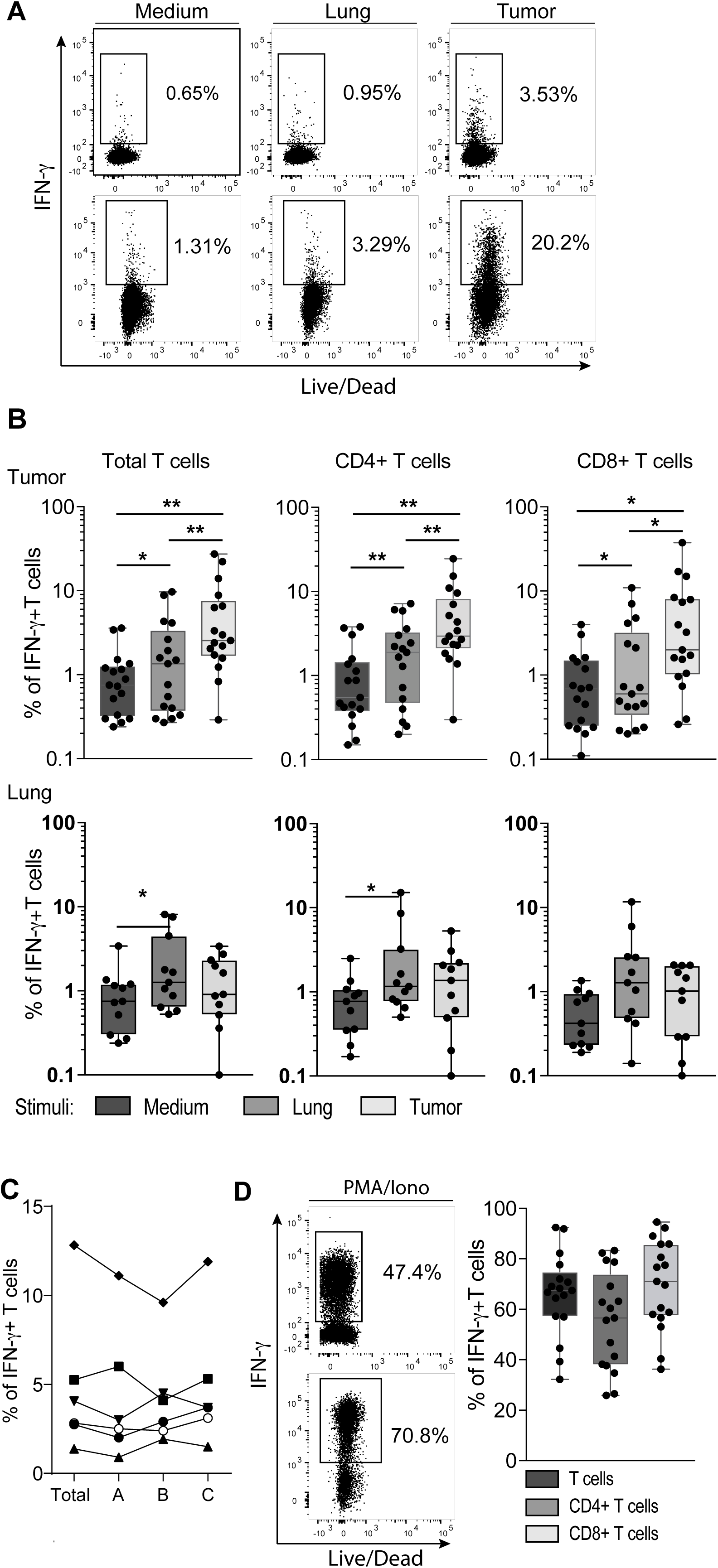
Most NSCLC-derived TIL products contain tumor-reactive T cells. **(A, B)** REP TILs were pre-stained with CD4 and CD8 antibodies, and cultured for 6 hours in the presence of brefeldin A with medium alone, with digests from normal lung tissue, or with autologous tumor digest. **(A)** Representative dot plots, and **(B)** compiled data of TIL products derived from 17 donors and (**C**) of T cells expanded from non-tumorous lung tissue from 11 donors of IFN-γ producing CD3^+^ T cells (left panel), by CD4^+^ T cells (middle panel) and by CD8^+^ T cells (right panel) as determined by flow cytometry. Each dot represents one donor, box and whisker plots depict minimum, 25^th^ pct, median, 75^th^ pct and maximum values. **(D)** IFN-γ production of CD4^+^ and CD8^+^ T cells compiled in response to tumor tissue from 6 donors extracted from **(B)**, of which three independent tumor regions (A-C) were isolated to compare the outgrowth of tumor-reactive TILs. (**E)** IFN-γ production by CD4^+^ and CD8^+^ T cells upon activation with PMA-Ionomycin. [Paired student’s T test; * p<0.05; ** p<0.01; *** p<0.001. If no indication, p≥0.05].

Interestingly, we detected 2.3±2.9% IFN-γ-producing T cells in the TIL products upon co-culture with lung digests, which was higher than those of TILs cultured with medium alone (1.1±1.0%; Fig. 2A, B). The reactivity could be due to undetected tumor cell dissemination, distal tumor antigen presentation by dendritic cells, bystander T cells that are not tumor-specific and thus may also respond to normal lung cells (24). The reactivity to normal lung tissue digests could also reflect innate immune responses by T cells that are triggered in a non-antigen specific manner (25, 26). We only considered TIL products tumor-reactive when the percentage of IFN-γ producing T cells was higher in response to the tumor digest than that to the normal lung digest. 13/17 (76%) of the TIL products fulfilled this requirement (see also below). We also tested the cytokine production of T cells that were expanded from lung digests. Only 2 out of 11 T cell products produced IFN-γ upon exposure to normal lung tissue, and only one lung-derived T cell product produced IFN-γ upon co-culture with tumor digest (Fig. 2B), indicating that the IFN-γ production from TILs is tumor-specific. Combined, these data indicate that tumor-reactive T cells are highly prevalent in expanded TIL products from NSCLC.

The percentage of T cells producing IFN_-_γ varied considerably between donors, ranging from 0.3% to 27.5% (Fig. 2B). This variation in tumor-reactivity could result from high heterogeneity of NSCLC tumors and thus depend on the tumor region that was isolated for TIL expansion. However, we consider this possibility unlikely, because the percentage of IFN_-_γ^+^ T cells expanded from whole tumor lysates was comparable to that of TILs that were expanded from three distinct parts of the same tumor (Fig. 2C). Furthermore, all TIL products produced substantial levels of IFN-γ in response to PMA/ionomycin stimulation (65.8±16.4%; Fig. 2D). There was no significant correlation between this potential to produce cytokines and the tumor-reactivity of TIL products (r=0.11, p=0.663). We therefore conclude that the majority of NSCLC-derived TIL products contain tumor-reactive T cells, but that the proportions thereof are variable.

### IFN-**γ**-producing tumor-reactive T cells express CD137 (4-1BB) and CD154 (CD40L)

We next interrogated if measuring the production of IFN_-_γ alone was sufficient to reflect the actual tumor-reactivity of TIL products. We therefore included the analysis of markers that are rapidly induced upon T cell activation, such as the costimulatory molecules CD137 (4-1BB) (27), and CD154 (CD40L) (28). When expanded TILs were co-cultured for 6h with tumor digest, we found a significant induction of CD137 expression in both CD4^+^ and CD8^+^ T cells (Fig. 3A, B, top panels). Furthermore, CD4^+^ T cells - but not CD8^+^ T cells - showed increased expression of CD154, a marker that was shown to identify antigen-specific CD4^+^ T cells in the peripheral blood (28) (Fig. 3A, B, middle panels). CD279 (PD-1) expression instead was not induced in expanded TILs within the 6h of co-culture with tumor digest (Fig. 3A, B, bottom panels), suggesting that the tumor-reactive T cells are not (immediately) susceptible to negative regulation by immunological checkpoints.

**Figure 3:**
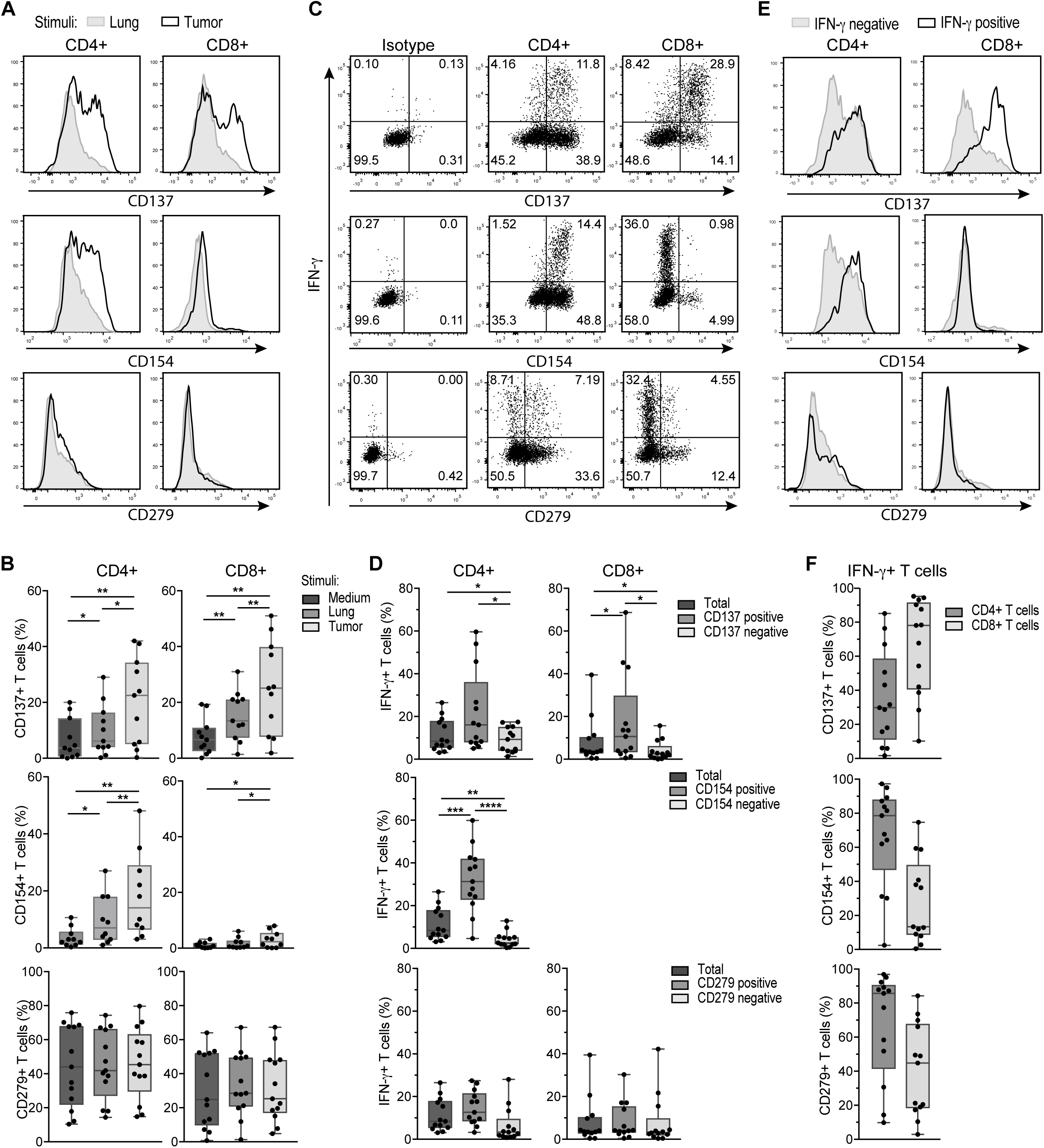
IFN-γ producing TILs express CD137 and CD40L, but not PD-1. REP TILs were cultured for 6 hours in the presence of brefeldin A with medium alone, with digests from normal lung tissue or with autologous tumor digest. **(A)** Representative histograms of CD137 (4-1BB), CD154 (CD40L), and CD279 (PD-1) expression of CD8^+^ and CD4^+^ TILs that were exposed to lung digest (gray), or to tumor digest (black line). **(B)** Compiled data of 13 TIL products for CD4^+^ T cells (left panel) and CD8^+^ T cells (right panel). Gates are set to isotype control. **(C)** Representative dot plots and **(D)** compiled data of percentage of T cells that co-express IFN-γ and CD137, CD154, or CD279 of CD4^+^ T cells (left panel) and CD8^+^ T cells (right panel). Gates are set to isotype control (C, left panels). **(E)** Representative histograms and **(F)** compiled data of the percentage of CD137, CD154, and CD279 –expressing CD4^+^ and CD8^+^ T cells within the IFN-γ producing (positive) and negative cells. Gates are set to isotype control. Each dot represents one donor, box and whisker plots depict minimum, 25^th^ pct, median, 75^th^ pct and maximum values. [Paired student’s T test; * p<0.05; ** p<0.01; *** p<0.001. If no indication, p≥0.05].

The majority of IFN-γ producing CD8^+^ T cells were CD137-positive (Fig. 3C,D). Combining the expression of CD137 and CD154 with the production of IFN-γ in fact helped to accurately define the presence of tumor-reactive TILs, in particular when the percentage of IFN-γ production was low, and/or the background levels high. However, we also observed that IFN-γ-negative T cells can express CD137 and CD154 (Fig. 3E,F), suggesting that the actual percentage of tumor-reactive T cells may be higher than estimated based on the production of IFN-γ.

### NSCLC-reactive TILs produce TNF-**α** and IL-2, and can be polyfunctional

In addition to IFN-γ, other cytokines such as TNF-α and IL-2 play a critical role in effective T cell mediated immunity (29, 30). This prompted us to determine whether TIL products from NSCLC produced these two cytokines. Interestingly, exposure to tumor digest resulted in significant induction of both TNF-α- and IL-2-producing T cells when compared to co-culture with lung digest or medium alone (Fig. 4A, B; Supplemental Fig. 2A). This was observed for both CD8^+^ T cells, and for CD4^+^ T cells (Fig. 4A, B).

**Figure 4:**
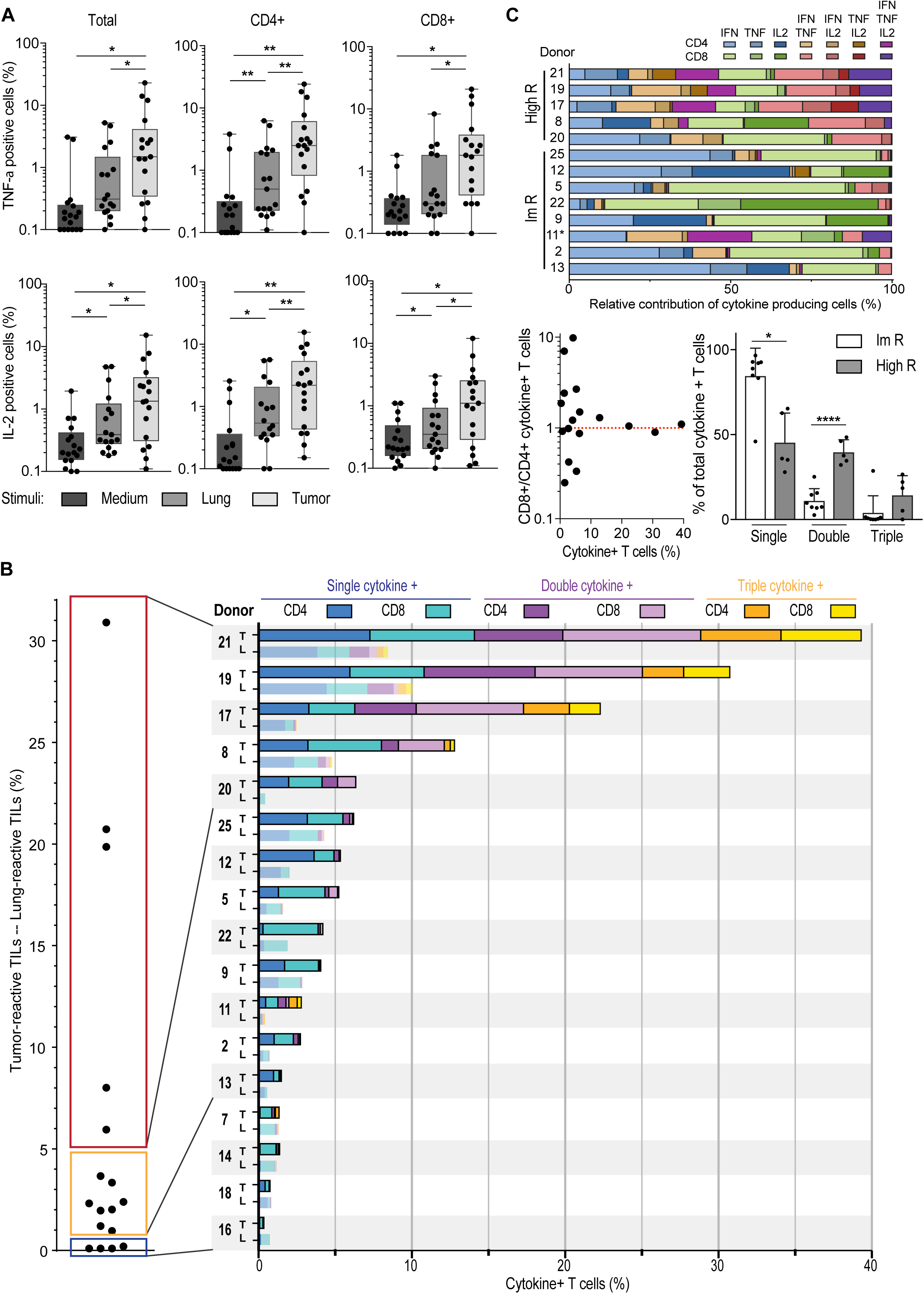
NSCLC-derived TIL products can be polyfunctional. **(A)** Percentage of TNF and IL-2 production of T cells (left panel), CD4^+^ T cells (middle panel) and CD8^+^ T cells (right panel) as determined by flow cytometry, n=17 donors. Each dot represents one donor, box and whisker plots depict minimum, 25^th^ pct, median, 75^th^ pct and maximum values. **(B)** *Left panel*: Percentage of cytokine-producing T cells that were cultured with tumor digest, minus the percentage of cytokine production of T cells stimulated with lung digest (n=17). *Right panel*: Total percentage of cytokine producing T cells. The percentage of cytokine producing CD4^+^ T cells (dark colors) and CD8^+^ T cells (light colors) is depicted when exposed to tumor digest (top bar), or to the lung digest (bottom bar). Color coding indicates the production of 1 (blue) 2 (purple), or 3 (yellow) cytokines simultaneously. Each patient is indicated with a number. **(C)** Top: Relative contribution of cytokine production profile of high and intermediate responders depicted in panel B. non-responders were excluded from this analysis. Bottom left: Ratio of tumor reactive CD8^+^ over CD4^+^ T cells, in relation to the percentage of cytokine-producing T cells. Bottom right: relative contribution of single, double and triple cytokine producers in high responders (high R), and intermediate responders (ImR) as defined in panel B.

When we used all three cytokines to define tumor-reactive TILs above background levels (i.e. subtracting the response against lung tissue), again 13/17 TIL products (76.5%) were tumor reactive (Fig. 4B, left panel). We next divided tumor-reactive TIL products into three categories. 4/17 (23.5%) expanded TILs did not produce cytokines above background levels. 8/17 (47.1%) TIL products contained up to 5% of cytokine-producing T cells (intermediate responder, Fig. 4B, left panel). Importantly, 5/17 (29.4%) TIL products contained at least 6%, and up to 31% cytokine-producing T cells above background levels (high responder; Fig. 4B, left panel). We also detected CD107 expression in the tested TIL products upon co-culture with tumor digest (Supplemental Fig. 4). Furthermore, high responders contained TIL products from SCC (patient 8, 17 and 19) and from AC (patient 20 and 21). Also, all tumor stages are equally distributed in high, intermediate and non-responders. These findings thus reveal that TIL products from NSCLC can contain high levels of tumor-reactive T cells.

Polyfunctional T cells that produce more than one cytokine simultaneously are considered the most potent effector T cells against chronic infections (31, 32). This is also observed for T cell responses against tumors (29, 30). We therefore investigated whether NSCLC-derived TIL products contain these polyfunctional T cells. Strikingly, all high responders contained T cells that produced 2 or 3 cytokines (Fig. 4B, right panel). Of the 8 intermediate responders, patient 7 and 11 also contained polyfunctional T cells (Fig. 4B, right panel). Polyfunctional responses to tumor digests were observed for both CD4^+^ and CD8^+^ T cells (Fig. 4B, right panel). Single cytokine producers included all three cytokines (Fig. 4C, top panel). Likewise, double cytokine producers could produce 2 out of the three cytokines in all possible combinations (Fig. 4C, top panel). Nevertheless, whereas intermediate responders (1-5% of total cytokine production; Fig 4B) were more skewed towards single cytokine producers, high responders contained single double and triple producers (Fig. 4C, bottom right panel). Interestingly, tumor-reactive T cells were also more equally distributed between CD4^+^ and CD8^+^ T cells in high responders (Fig. 4B, right panel), and the overall ratio of CD8^+^ T cells over CD4^+^ T cells in REP products of high responders was close to 1, whereas this ratio was not skewed to either T cell subset in intermediate and no responders (Fig. 4C, bottom left panel). In conclusion, most TIL products contained tumor-reactive CD4^+^ and CD8^+^ T cells, and in particular high responders comprised of polyfunctional T cells, indicative for potent anti-tumoral TIL products generated from NSCLC tumors.

### Tumor-specific alterations in T cell composition correlate with tumor-reactivity of TIL products

We next investigated whether the high variation in tumor reactivity between TIL products related to specific features of T cells in the original tumor tissue. Because the composition of T cell infiltrates is highly variable between individuals (see below), we used the normal lung tissue as base line for each patient to define the tumor-specific T cell signature.

Tumor tissues showed a modest, but significant increase in CD8^+^ T cell infiltrates compared to lung tissue, a feature that was not observed for CD4^+^ T cells (Fig. 5A). Tumor lesions contained in particular higher percentages of CD27^+^ CD45RA^−^ central memory T cells (Supplemental Fig. 5), as well as FOXP3^+^CD25^high^ CD4^+^ T cells (Fig. 5B). These FOXP3^+^ CD4^+^ T cells did not express the IL-7 receptor alpha chain (CD127), identifying them as bona fide regulatory T cells (33) (Fig. 5B).

**Figure 5:**
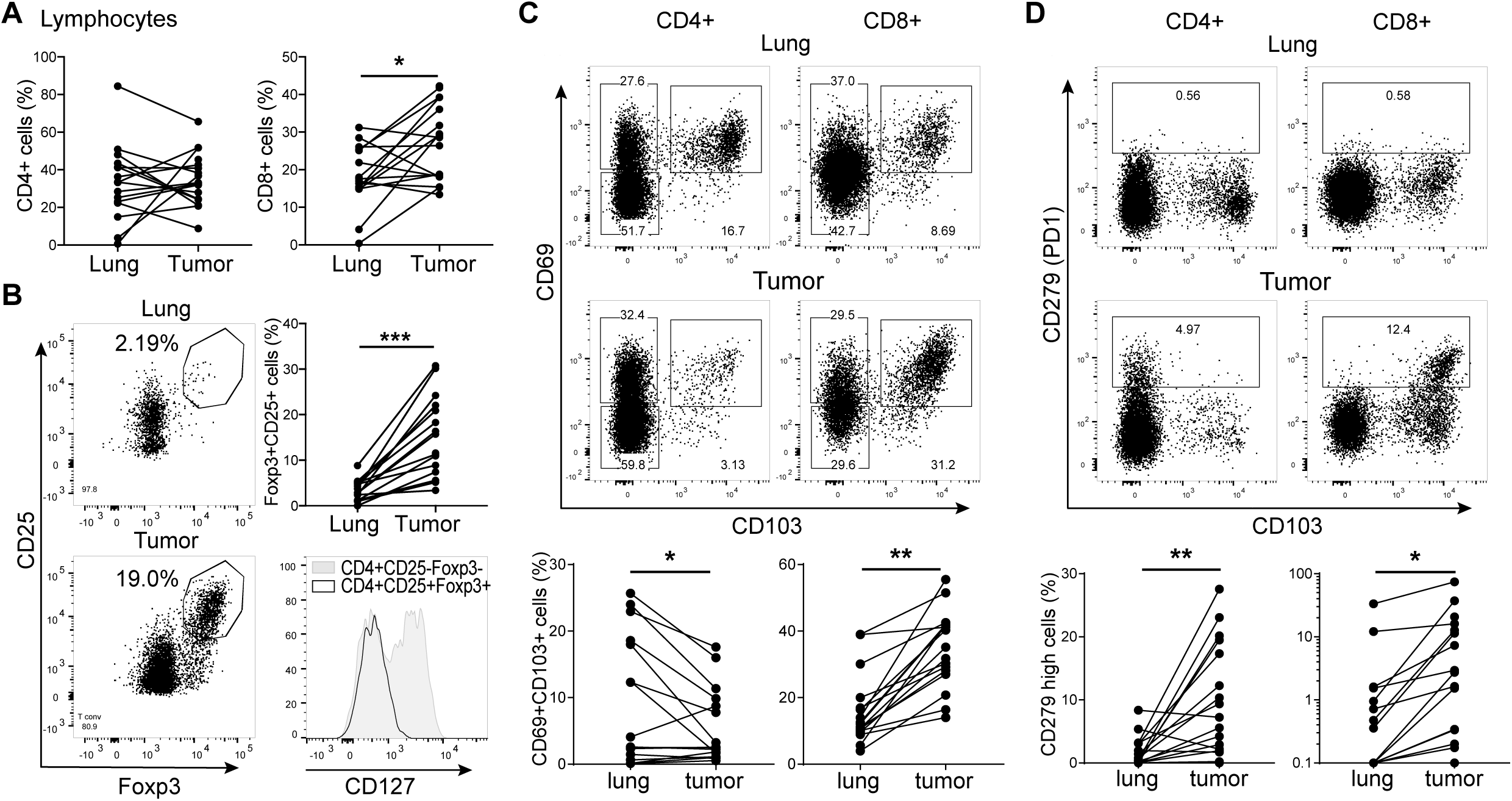
T cell composition is altered in the tumor lesions. **(A-D)** Tumor tissue and of normal lung tissue digests were analysed directly *ex vivo* by flow cytometry to determine the expression of CD3^+^CD4^+^ (left panels) and of CD3^+^CD8^+^ T cells (right panels) within the lymphocyte population **(A)**, of CD4^+^CD25^+^ Foxp3^+^ T cells **(B)**, of CD69^+^CD103^+^ T cells **(C)** and of CD279^+^ T cells **(D).** Data are shown as representative dot plots and compiled data of all 17 patients. In **(B)** a representative histogram is shown of CD127 expression on tumor derived CD25^+^Foxp3^+^CD4^+^ T cells versus conventional FoxP3^−^CD4^+^ T cells. [Paired student’s T test; * p<0.05; ** p<0.01; *** p<0.001. If no indication, p≥0.05].

The percentages of CD8^+^ T cells that express both the integrin CD103 and the retention marker CD69 in the tumor lesion significantly exceeded those in normal lung, which were already high (Fig. 5C). For CD4^+^ T cells, we instead observed a drop of CD69^+^CD103^+^ expressing cells in the tumor when compared to the normal lung tissue (Fig. 5C). As expected, we found that PD-1^hi^ expressing CD4^+^ and CD8^+^ T cells were substantially increased in the tumor lesions compared to normal lung tissue (Fig. 5D).

Thus, tumor tissues contain high levels of FOXP3^+^CD25^high^ regulatory CD4^+^ T cells, PD-1^hi^ CD4^+^ and CD8^+^ T cells, and high levels of CD69^+^CD103^+^CD8^+^ T cells.

We next investigated whether a specific *ex vivo* tumor-specific T cell signature correlated with the level of tumor-reactivity detected in the TIL products. To account for the high variation between patients already in the healthy lung (Fig. 5), we used the tumor-specific increase of T cell subpopulations for this comparison (delta percentage; Δ%). The tumor-specific infiltration by CD4^+^ or CD8^+^ T cells, or the effector/memory phenotype of both T cell subsets did not correlate with the percentage of cytokine producing TILs (Fig. 6A, Supplemental Fig. 5). Also the expression levels of PD-L1 on tumor cells or on immune infiltrates did not indicate high tumor reactivity (Fig. 6B). However, when examining specific T cell subsets, we found that the percentage of CD69^+^CD103^+^ CD8^+^ T cells positively correlated with the level of tumor-reactivity of expanded TIL products (Fig. 6B). For CD4^+^ T cells, a positive correlation with tumor-reactivity was found for the percentage of PD-1^hi^ CD4^+^ T cells (Fig. 6B,C). Intriguingly, even though regulatory T cells are associated with a poor prognosis for NSCLC patients (17, 18), we found that the presence of tumor-specific FOXP3^+^CD25^hi^ CD4^+^ T cell infiltrates correlated positively with cytokine production of expanded TILs (Fig. 6D). Combined, our data suggest that the tumor-specific signature of CD4^+^ and CD8^+^ T cells and a high number of Treg infiltrates is indicative for strong tumor reactivity in the expanded TIL products.

**Figure 6:** Tumor-specific T cell infiltrates correlate with cytokine production of the TIL product. Pearson’s correlation of the percentage of cytokine production of expanded T cells as defined in Fig. 4 with tumor-specific infiltrates of **(A)** CD4^+^ (left panel) CD8^+^ (right panel) CD3^+^ T cells, **(B)** PD-L1 expression on tumor cells and on immune cells, **(C)** CD69^+^/CD103^+^ T cell population within CD4^+^ (left) and CD8^+^ (right) T cells, **(D)** with CD279^hi^ CD4^+^ T cells (left panel) and CD8^+^ T cells (right panel), and with **(E)** with Foxp3^+^/CD25^high^ CD4^+^ T cells. (** p<0.01;*** p<0.001. If no indication, p≥0.05).

## DISCUSSION

Here we show that tumor-specific TILs can be effectively expanded from the majority of NSCLC patients. Importantly, we found that TIL products that contain high percentages of tumor-reactive T cells produce not only IFN-γ, but also TNF-α and/or IL-2. T cells producing more than one cytokine are considered highly potent (29, 30) and vaccination strategies strive to generate these polyfunctional T cell responses (34). We therefore hypothesize that infusion of TIL products containing polyfunctional T cells should be effective against NSCLC.

Recently, it was shown that tumor-reactive TILs could be expanded from 80% of tested melanoma patients that did not respond to anti-PD-1 treatment (35). 2 out of 12 patients that were subsequently treated with TIL therapy showed objective clinical effects (35). These findings thus indicate that melanoma patients who are refractory to anti-PD-1 can still benefit from TIL therapy, and therefore anti-PD-1 and TIL therapy are complementary immunotherapeutic strategies. This could also be the case for NSCLC patients that did not respond to anti-PD-1 treatment (9–13).

Intriguingly, most NSCLC TIL products contain both tumor-reactive CD8^+^ T cells and CD4^+^ T cells. The cytolytic function of CD8^+^ T cells to control and/or eradicate tumors is well established, and tumor-reactive CD8^+^ T cells were recently expanded by others from NSCLC tumor digests and from peripheral blood (23, 36, 37). In our study the majority of TIL products contained tumor-reactive CD8^+^ T cells.

Much less is known about tumor-reactive CD4^+^ T cells. Recently, neo-antigen specific CD4^+^ T cells were identified in the majority of tested melanoma lesions (38). Furthermore, two patients that suffered melanoma and metastatic cholangiocarcinoma received TIL therapy with tumor-reactive CD4^+^ T cells only, and both patients showed clear clinical benefits from this treatment (39, 40). Thus, CD4^+^ T cells can substantially contribute to anti-tumoral responses. Whether this effect is indirect through promoting effective CD8^+^ T cell responses, or direct by cytotoxic CD4^+^ T cells as proposed in murine tumor models (41, 42) is yet to be defined. These findings combined strongly suggest that in addition to the CD8^+^ T cells, these tumor-reactive polyfunctional CD4^+^ T cells could be of value for TIL therapy.

The composition of the tumor-infiltrating T cell compartment correlates with patient survival (43–46). Our study now showed that a correlation also exists between this composition and the likelihood that cytokine-producing tumor-reactive T cells are present in expanded TIL products. In line with previous studies that showed that the presence of CD8^+^ T cells expressing CD103 and CD69 serves as a favorable prognostic marker for survival (43, 44, 47), we describe here that their presence in the tumor lesion also correlated with high percentages of cytokine-producing CD8^+^ T cells in the TIL products. This was, however not the case for CD4^+^ T cells, for which high PD-1 expression correlated with the presence of tumor-reactive T cells in the TIL product.

To our surprise, even though the presence of FOXP3^+^ T cells in tumors is associated with poor survival (45, 46), we found the highest levels of tumor-reactivity in TIL products when the percentage of Tregs in the tumor lesion was also high. One possible explanation could be that Tregs are recruited to quench ongoing T cell responses against the tumor, such that their prevalence would be proportional to the magnitude of the anti-tumor response (48). Irrespective of the mechanism(s) that contribute to the correlation of high Treg numbers with high percentages of tumor-reactive T cells, it is tempting to speculate that patients with high Treg numbers may profit most from TIL therapy. Furthermore, even though a validation cohort of our findings is warranted, we propose that the presence of CD69^+^CD103^+^ on CD8^+^ T cells, PD-1^hi^ CD4^+^ T cells, and high levels of Tregs in the tumor lesions may help identify the patients that are most likely to benefit from TIL therapy.

In conclusion, we here demonstrate that most tumor-reactive TIL products generated from NSCLC tumor lesions contain tumor-reactive T cells. We therefore suggest that TIL therapy should be considered as treatment for NSCLC patients.

## DECLARATION STATEMENTS

### Ethics approval and consent to participate

The study was performed according to the Declaration of Helsinki (seventh revision, 2013), and executed with consent of the Institutional Review Board of the Netherlands Cancer Institute-Antoni van Leeuwenhoek Hospital (NKI-AvL), Amsterdam, the Netherlands (#CFMPB317).

### Consent for publication

Not applicable.

### Availability of data and material

All data generated or analysed during this study are included in this published article [and its supplementary information files].

### Authors’ contributions

RdG, MMvL, MMvdH, JdJ, PB, CEvdS, RMS, DA, JBAGH, KM, KJH and MCW designed the study; RdG, MMv, AG, BPN, JJFvH, KM, and OV performed experiments and analysed data, KJH coordinated sample collection and performed pulmonary surgery, JdJ, KM collected and analysed samples. RdG and MCW wrote the manuscript. MCW supervised the study. All authors read and approved the final manuscript.

## Acknowledgements

We thank the Flow cytometry facility from Sanquin Research, and the medical assistance staff from the NCI-AvL for technical help.

---

**Supplemental Figure 1** Top panel: Pearson’s correlation of fold T cell expansion as defined in Fig. 1 with PD-L1 expression on tumor cells and on immune cells as defined by immunohistochemistry (Table 2). Middle panels: Pearson’s correlation of CD4^+^ T cells and bottom panels: CD8^+^ T cells with tumor-specific T cell infiltrates, as defined in Supplemental Fig. 5.

**Supplemental Figure 2 (A)** Percentage of CD25^hi^CD127^−^FOXP3^+^ CD4^+^ Treg cells ex vivo, and upon expansion. n=4 donors. **(B).** Representative histograms, and percentage of T cells expressing PD-1, LAG-3, and TIGIT ex vivo, and after expansion, as determined by flow cytometry. n=4 donors.

**Supplemental Figure 3:** T cell clone count (log2) from tumor lesions compared to expanded TILs based on the nucleic acid sequence of the CDR3 region of the TCRbeta chain. Top row: CD4^+^ T cells, excluding the CD25^hi^CD127^hi^ CD4^+^ cells containing the Treg population. Bottom row: CD8^+^ T cells. Each plot depicts one donor.

**Supplemental Figure 4 (A)** REP TILs were pre-stained with CD4 and CD8 antibodies, and cultured for 6 hours in the presence of brefeldin A with medium alone, with digests from normal lung tissue, or with autologous tumor digest or PMA/Ionomycine. Representative dot plots of the production of TNF-α and IL-2 of one high and one intermediate donor (same as in Fig. 2A) by total T cells as determined by flow cytometry. **B)** CD107 expression of expanded TILs after 6h incubation with medium (left), lung digest (middle) and with tumor digest.

**Supplemental Figure 5** Left panels: representative dot plots of the effector/memory phenotype of CD4^+^ T cells CD8^+^ T cells from tumor digest based on CD45RA and CD27 expression. Right panels, 1^st^ and 3^rd^ row: Percentage of naïve (TN), central memory (TCM), effector memory (TEM) and CD45RA^+^ effector memory (TEMRA) T cells in the lung, and in the tumor. Right panels, 2^nd^ and 4^th^ row: Pearson’s correlation of the percentage of cytokine production of expanded T cells as defined in Fig. 4 with tumor-specific effector/memory T cells infiltrates.

